# Occurrence of Biased mTOR Signaling in Hepatocellular Carcinoma

**DOI:** 10.64898/2026.05.22.727188

**Authors:** Rudransh Singh, Niriksha Patel, Neha Singh, Pooja Mourya, Ashish Shingade, Anjali Mange, Jasmine Kaur, Sakshi Beloshe, Ashok Dudhalkar, Pruthviraj Chavan, Ghanapriya Devi Yengkhom, Shraddha Patkar, Mahesh Goel, Arvind Ingle, Sagar Ranjan Tripathy, Sridhar Epari, Sharathchandra Arandkar, Rahul Thorat, Sunil Shetty

## Abstract

**Background:** mTOR signaling promotes cell growth and anabolic processes in all eukaryotes. Hyperactivation of mTOR signaling is associated with various cancers along with hepatocellular carcinoma (HCC). HCC is a highly lethal malignancy with multiple aetiologies such as viral infection, alcohol abuse, and metabolic dysfunction. Therapeutic options for HCC remain limited due to an incomplete understanding of oncogenic drivers and poorly characterized mechanisms of disease progression.

**Methods:** Various regimens of DEN and CCl_4_ carcinogen dosage were investigated on C57BL/6J mice to induce HCC. The histological analysis for fibrosis and serum markers for liver function were performed. Molecular analyses of oncogenic drivers were performed in the HCC tissues obtained from mice and HCC patients. The impact of inhibition of mTOR signaling was assessed on HCC progression.

**Results:** We established a rapid DEN+CCl_4_ induced (DCI) HCC model in C57BL/6 mice to study disease progression longitudinally. The molecular analysis revealed upregulation of MAPK and downregulation of mTORC1-S6K-S6 signaling in HCC. However, other branches of mTOR such as mTORC1-ULK1, mTORC1-4EBP1, and mTORC2-PKCα were upregulated due to the increased expression. Similar observations were found in tissues derived from HCC patients. Furthermore, inhibition of mTORC1 alone by Rapamycin did not reduce HCC progression but Torin 1 mediated inhibition of both mTORC1 and mTORC2 significantly reduced HCC progression.

**Conclusions:** We propose this biased mTOR signaling modulates mTOR activity towards specific downstream processes that are crucial for cancer cell growth and targeting both the mTOR complexes has better therapeutic potential in HCC.

**Impact and Implications:** This study provides a rapid pre-clinical model for understanding HCC progression and to explore various intervention strategies. The study reports a novel phenomenon of biased mTOR signaling where deregulation of downstream substrate levels modulates the mTOR activity towards the specific branches, the master regulator of cell growth and metabolism. Furthermore, the study suggests that the clinical investigations exploring the rapalogs against HCC should be cautiously considered depending on the aetiology and signaling status of HCC.

**Graphical abstract:** 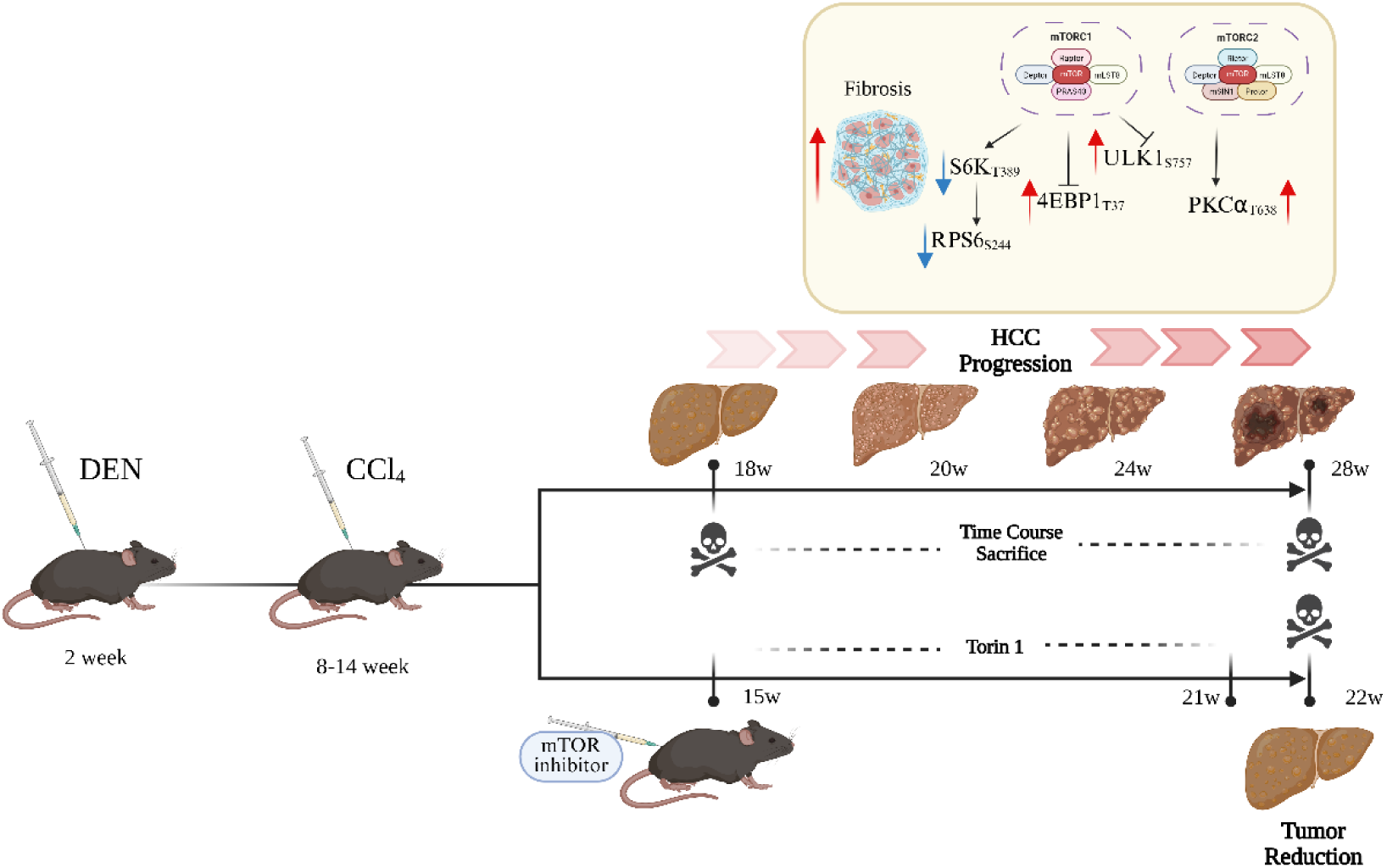

**Highlights:**

- DEN and CCl_4_ treatment generated a well-established HCC within 4 months.
- Liver fibrosis and serum markers correlated with HCC progression.
- Upregulation of mTOR pathway substrates create biased mTOR signaling.
- Dual inhibition of mTORC1 and mTORC2 reduced HCC progression significantly.

## Introduction

Hepatocellular carcinoma (HCC) is an aggressive malignancy marked by rapid progression and poor prognosis. The occurrence of disease ranks 8^th^ among all cancer types and 4^th^ in cancer related mortality, suggesting its highly fatal nature (1). The disease typically develops through a multistep process starting from the chronic liver injury, leading to inflammation and hepatocellular degeneration (2, 3). This is followed by the emergence of dysplastic nodules, finally demonstrating HCC. Surgical intervention as a therapeutic option is often limited to early stage patients (2, 3). As most patients are diagnosed at the late stages with symptoms like advanced liver fibrosis or even cirrhosis, the patients are often offered targeted therapies such as Lenvatinib or Sorafenib (4–7). Recently approved immunotherapies, including the anti-PD-L1 antibody Atezolizumab (often combined with other agents), benefit only selected patient subgroups (8). The success of targeted therapy using multi-kinase inhibitors is often limited with emergence of resistance by pathways such as mTOR (9).

One of the major hurdles in understanding the molecular drivers is the diversity in the aetiology of HCC. HCC can be caused by Hepatitis B or Hepatitis C viral infection, excessive alcohol consumption, aflatoxin, and metabolic dysfunction. The surge in cases of obesity and fatty liver diseases are increasing the burden of metabolic-dysfunction associated HCC cases (10). These factors trigger chronic inflammation along with genetic and epigenetic alterations that promote hepatocarcinogenesis. These multifactorial stimuli also contribute to disease complexity and limited therapeutic success (11). Thus, better understanding of the aetiology specific oncogenic molecular drivers is necessary for developing improved targeted approaches.

*In vivo* pre-clinical animal models are instrumental in understanding the aetiology specific molecular drivers, as HCC can be induced with the treatment of chemicals, alcohol or dietary changes in the immune-competent rodents(12–14). The chemical carcinogen-induced rodent models being particularly relevant as they recapitulate HCC associated with metabolic and alcohol-related aetiologies(15). A commonly employed approach is the two-stage model(16), where diethylnitrosamine (DEN) initiation is followed by chronic exposure to genotoxic or non-genotoxic promoters (e.g., 2-AAF, NMOR, phenobarbital, ethanol, CCl₄)(15, 17). DEN induces persistent genetic mutations in proliferating hepatocytes(18, 19), while carbon tetrachloride (CCl₄) generates reactive free radicals that promote lipid peroxidation and membrane damage(20, 21). This oxidative stress activates Kupffer cells and triggers inflammatory cascades involving cytokines and chemokines, leading to immune cell recruitment and amplified hepatic injury(22–24). This regimen mimics initiation–promotion processes, inducing preneoplastic lesions, fibrosis, and HCC in rodents. This repeated cycles of injury, inflammation, and regeneration ultimately drive fibrosis and hepatocarcinogenesis, and enabling mechanistic studies of tumor progression(23). Despite its strong hepatocarcinogenic potential, the CCl₄-based model requires prolonged exposure (≈6–12 months) for tumor development(25, 26), resulting in extended study durations and high animal mortality(23).

Proteogenomic analyses of hepatocellular carcinoma (HCC) patients have revealed substantial molecular heterogeneity in tumor drivers, accompanied by the activation of multiple growth-promoting signaling pathways, including Wnt/β-catenin, MAPK, and PI3K/AKT/mTOR(27, 28). Approximately 50–60% of HCC cases exhibit dysregulation of Wnt/β-catenin and MAPK signaling, depending on disease stage(29), while nearly 40–50% show elevated mTOR pathway activity(30, 31). mTOR signaling is highly conserved eukaryotic kinase which exists as two complexes called mTORC1 and mTORC2 (32, 33). mTORC1 is the master regulator of cell growth and regulates diverse processes in a cell such as ribosome biogenesis, translation, autophagy, and transcription via its downstream mediators like S6K1, 4EBP1, ULK1, and TFEB (34–36). The mTORC2 signaling regulates cytoskeleton and cell survival pathways via AKT, SGK, and PKC kinases (37, 38). The activity and role of both mTORC1 and mTORC2 in the diverse aetiology of HCC remains obscure.

In this study, we established a mice model to study hepatocellular carcinoma (HCC) progression in C57BL/6 mice called DCI model using DEN and CCl₄ carcinogens. This study shows that a single dose of DEN and six weeks of CCl_4_ treatment can induce HCC progressively within 6-10 weeks. This model can be used for the longitudinal investigation of disease progression. Molecular analyses of HCC tissues revealed activation of MEK–ERK and Wnt signaling pathways. Interestingly, mTOR signaling showed varied activity depending on downstream substrates. We find reduced activity of mTORC1-S6K-S6 axis while upregulation of mTORC1-4EBP1, mTORC1-ULK1 and mTORC2-PKCα axis in the DCI model. We propose this phenomenon as elicitation of biased mTOR signaling which is mainly driven by the upregulation of total levels of specific downstream substrates of mTOR signaling. The analysis of patient-derived HCC-tissues also shows the elicitation of biased mTOR signaling in some patients. Furthermore, we observed that inhibition of both mTORC1 and mTORC2 by Torin 1 had better therapeutic potential than inhibition of mTORC1 alone by Rapamycin.

## Materials and Methods

### Ethical approval

The study was conducted in accordance with the guidelines approved by the Institutional Review Board (IRB) and Ethics Committee (EC) of the Tata Memorial Centre–ACTREC (IEC No. 901029). Patient samples were obtained from the tumor tissue repository (TTR) at Tata Memorial Hospital (TMH) and the biorepository at the Advanced Centre for Treatment Research and Education in Cancer (ACTREC), Mumbai, which maintain fresh-frozen tumor and matched normal tissues for research purposes. As samples were collected retrospectively, the requirement for informed consent was waived by the IRB and EC, in line with standard institutional procedures at TMH and ACTREC. Also, for the animal studies, the project was approved by the Institutional Animal Ethics Committee (IAEC) and all the experiments were conducted according to the guidelines directed by the committee (Project No. 25.2023).

### Mouse Model establishment and tissue collection

The C57BL/6J mice were kept under a standard light/dark cycle of 12 hours, and had access of the food and water ad libitum. The male mice were divided in multiple groups according to the treatment regimen for the carcinogen dosing. The C57BL/6J male mice under the treatment group were injected with single dose of DEN 25mg/kg intraperitonially at 2^nd^ week of age and the control group mice received 0.9% saline. From the age of 8^th^ week, the DEN administered mice were treated with CCl_4_ (8ml/kg, 10% solution in olive oil) twice a week, for 6, 10 or 14 weeks as mentioned in the respective figures. The control cohort was treated with olive oil for the similar duration. The mice were humanely euthanized as per the schema shown in figure and liver tissue chunks were collected and snap frozen into liquid nitrogen. For the histology analysis, tissues were fixed in formalin solution. During the whole tenure of the study, the mice weight was measured at regular weekly intervals and also a day before sacrifice. For the increased DEN dose model, C57BL/6J mice were treated with 20, 40, and 60mg/kg of DEN at the age of 2^nd^, 4^th^, and 6^th^ weeks respectively.

### Histological examination, Reticulin, and Collagen Staining

Tissues fixed in formalin solution were embedded in paraffin blocks and their sectioning was done. Hematoxylin and Eosin (H&E) staining was performed as per the standard protocols. The reticulin staining was performed as per the previous Gomori’s reticulin protocol (39). For picrosirius red staining, the slides were deparaffinized via heating in hot air oven at 55°C for 20 min, followed by rehydration in a series of solvents i;e. xylene, xylene+ethanol (1:1), ethanol, 70% ethanol, and PBS. Then slides were incubated in hematoxylin solution for 5 minutes. After rinsing the slides in running tap water for 10 min, slides were immersed in Picrosirius red stain for 1 hour. Slides were rinsed with 0.5% glacial acetic acid briefly and dehydrated through solvents in reversed order (40). Slides were air-dried and coverslip was mounted using DPX mountant. The slides were scanned through TissueScop iQ slide scanner (Huran technologies) and the images were analysed with Qupath (Ver. 0.5.0). The images were further processed with ImageJ tool for intensity quantification.

### Chemicals and drugs treatment

The mice were treated with carcinogens to develop 6w-DCI model. This was followed by the intraperitoneal administration with Sorafenib (2mg/kg), Rapamycin (10mg/kg), and Torin 1(10mg/kg) from the age of 15 to 21 weeks. At the onset of each week before the drug injections, body weights were collected group wise and dosages were given thrice per week for 6 weeks. Animals were humanely euthanized at the age of 22 weeks.

### Serum analysis

After euthanization, the blood was collected through the cardiac puncture method. The blood samples were centrifuged gently at 3000 RPM at room temperature and serum was collected and stored in - 80°C, till further analysis. The liver function assays for various parameters such as total protein, albumin, cholesterol, bilirubin, triglycerides, alanine transaminase, alkaline phosphatase, and aspartate transaminase in serum samples were performed via an automated system using Atellica Solution (Siemens Healthineers).

### Protein extraction and immunoblotting

The mice liver tissue chunks were lysed in RIPA buffer (50mM Tris-Cl pH 7.4, 150mM NaCl, 1% triton X-100, 0.5% Sodium Deoxycholate, 1% SDS, 1mM EDTA) containing Protease inhibitors and phosphatase inhibitors. The tissues were homogenized using bead rupture method at 4°C (60sec pulse for 3 cycles with 60sec break). Then homogenates were collected and the protein content was measured by BCA assay. The total proteins (15ug per sample) were separated on 10 % SDS-PAGE gel and transferred onto nitrocellulose membranes. The membranes were processed for immunoblotting as per standard procedure and the blots were visualized using ECL reagent. The details of antibodies and dilutions are provided in supplementary table 1.

### RNA isolation from tissues and qPCR analysis

Total RNA was extracted from the tissue samples using the TRIzol reagent. RNA concentration was measured using the nanodrop spectrophotometer. First strand cDNA synthesis was performed using Primescript 1^st^ strand cDNA synthesis kit (Takara) as per the protocol. Quantitative real-time PCR was performed using Takara TBgreen II real time PCR master mix (Takara) as per the manufacturer’s instructions. qPCR was performed using conditions of initial denaturation at 95°C followed by 40 cycles of 95°C for 15s, 58°C for 18s, and 72°C for 15s. The mRNA expression was calculated using ΔΔCt method. List of all primers is provided in supplementary table 2.

### TCGA-LIHC data survival analysis

The RNA-seq data of LIHC patients was acquired from the TCGA-LIHC project (n=365). Additionally, patient’s survival information was procured from University of California Santa Cruz – XENA portal for the same set of patients. The survival plots were made using the survival and survminer tool in the R studio. The curated RNA seq data of Pan-cancer patients was also acquired form TCGA.

### Statistics and reproducibility

Statistical analysis was performed using GraphPad Prism version 8 (GraphPad Software, La Jolla, CA, USA). One-way ANOVA was used to determine the statistical significance between the two treated groups vs. control group, and the *p* values calculated are denoted above the significance mark in the plot. The reproducibility of the experimental findings was confirmed by performing *n* = 3 independent replicates. The results of all the biological replicates were consistent.

## Results

### Establishment of the DEN and CCl_4_-induced (DCI) mouse model

The major hurdle in establishing HCC animal models is the prolonged duration (6 to 12 months) for tumor development (12, 13, 15, 17, 25). As per the previous protocol, to develop HCC in mice, we administered C57BL/6 male mice with a single dose of DEN (25mg/kg) at the 2^nd^ week from birth followed by 14 weeks of CCl_4_ treatment (10% CCl_4_ in olive oil, 8ml/kg twice/week (16) (modified from the reference(26)) starting from the age of 8^th^ week (Fig. S1A(i)). In corroboration with previous study, we also observed robust development of HCC tumors when mice were sacrificed at the age of 28^th^ week, post 6 weeks of complete CCl_4_ regimen (Fig. S1A(ii) and (iii)). There was no significant difference in the body weight of mice between control and carcinogen treated group (Fig. S1A(iv)). Since this model showed extensive nodules in the entire liver, analysis of tumor progression was not feasible. To study HCC tumor progression, we reduced the number of doses of CCl_4_ treatment for 6- or 10-weeks, instead of 14 weeks. Thus, after the single DEN injection at the 2^nd^ week, the mice were treated with CCl₄ (10% CCl_4_ 8ml/kg twice/week) for either 6- or 10-weeks duration (a total of 12- or 20- CCl_4_ injections, respectively) from 8^th^ week (Fig. 1A (ii) and (iii)). Henceforth in this study, this model is referred as DCI model to indicate DEN-CCl_4_ induced HCC model with different duration of CCl_4_ regimen as prefix (6w-DCI, 10w-DCI, 14w-DCI). The control group received 0.9% saline against DEN, followed by olive oil instead of CCl₄ (Fig. 1A(i)). To capture the early events of tumorigenesis, sequential sacrifice has been performed for all the groups, and their body weight and liver weights were recorded. Although the body weight (Fig. S1B) was almost similar among different groups at each time point, there were significant difference in the liver weight (Fig. 1B) immediately post treatment with 10w-CCl_4_ (10w-DCI). In the case of 6w-CCl_4_ cohort (6w-DCI), there was no significant increase in the liver weight immediately post CCl_4_ treatment. However, post 10 weeks of CCl_4_ treatment (at the age of 24w), liver weight increased significantly, suggestive of hepatomegaly.

**Fig. 1.**
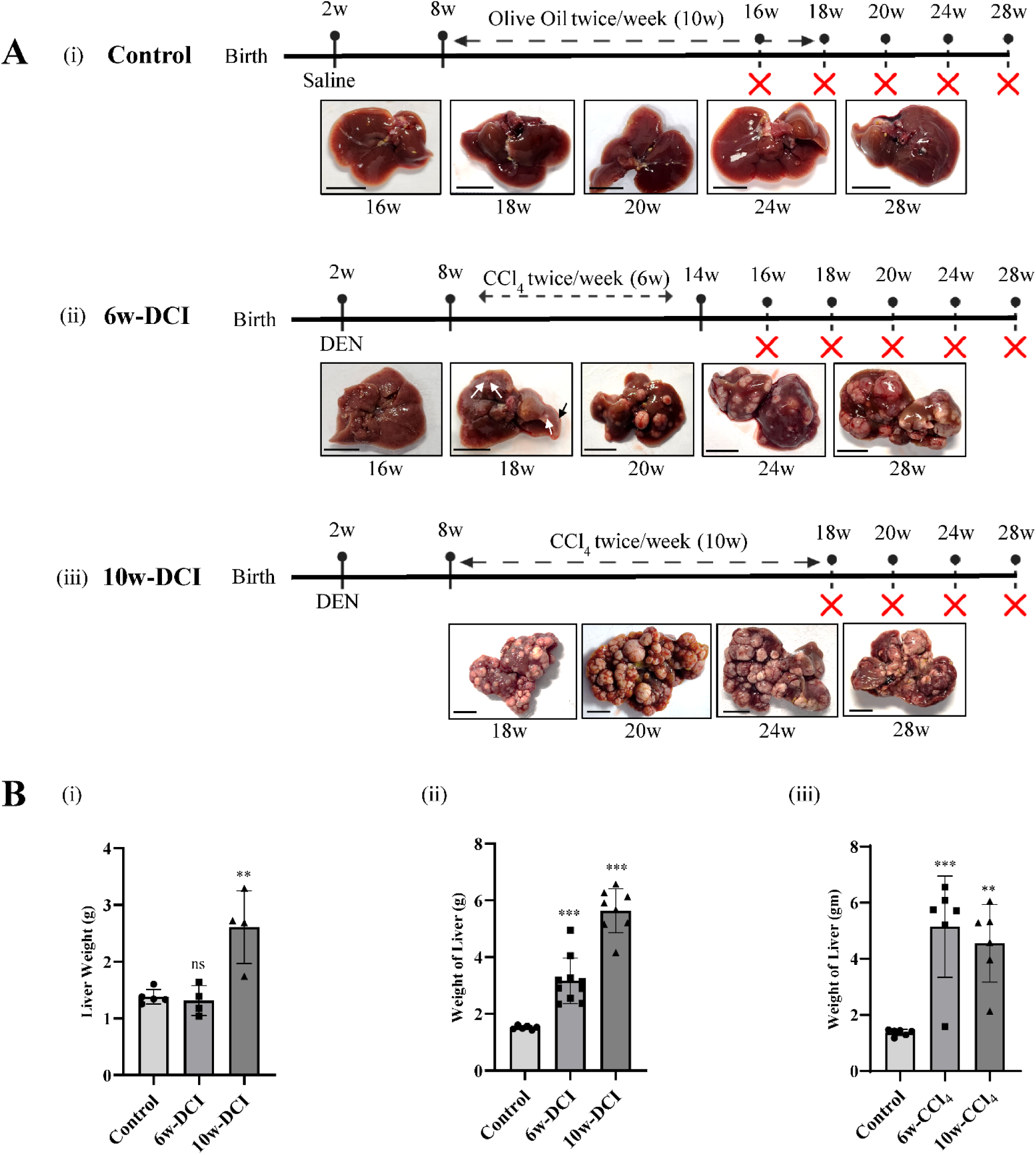
Establishment of DEN+CCl_4_-induced (DCI) HCC model in C57BL/6J mice. (**A**) The schematic of DEN and CCl_4_ or vehicle injection regimen and representative liver images upon sacrifice at various time points (n=4-6 per time point). (i), Control group is treated with saline at 2 weeks of age and olive oil from 8 to 18 weeks; (ii) 6w-DCI cohort is treated with DEN at 2 weeks of age and CCl_4_ from 8 to 14 weeks of age; (iii), 10w-DCI cohort is treated with DEN at 2 weeks and CCl_4_ from 8 to 18 weeks (Scale Bar = 1cm). The mice were sacrificed at 16, 18, 20, 24 and 28 weeks of age (**B**) Liver weight quantification after sacrifice of control, 6w-DCI and 10w-DCI at 18, 24, and 28 weeks. Data is represented as means ± SEM, plotted as bar graph. Y-axis represents weight values (g). Each dot represents liver weight of single mouse. *p-values* are from one-way ANOVA tests and, significance marks are indicated above respective bar in comparison to control. *p*-values are denoted as *ns p*>0.05; \*\**p* < 0.01; \*\*\**p* ≤ 0.001.

The 10w-DCI group possessed multiple HCC nodules at the age of 18 weeks, immediately post CCl_4_ regimen (Fig. 1A(iii)) in all mice (n=4). At the 18^th^ week, the nodule sizes were relatively smaller than the nodules observed in 14w-DCI model and also had non-tumor regions for comparative biochemical analysis. Further monitoring the mice for 4 to 8 weeks without CCl_4_ treatment showed significant increase in the nodule size, suggesting that the 10w-DCI model recapitulates the intermediate to late stage of HCC with the window of progressive tumor growth. Thus, we report a rapid method for chemical-carcinogen induced HCC model development within 4 months, significantly reducing the time required to generate HCC model compared to previous studies (13, 17, 25). In contrast, the 6w-DCI model (Fig. 1A(ii)) did not show any macroscopic tumors immediately (at the age of 16 weeks). However, the 6w-DCI mice showed slow development of HCC tumors post 6 weeks (n=4) of treatment while post 10 weeks showed clear visible tumors (at the age of 24 weeks, n=8). Thus, this 6w-DCI model serves as an ideal model to study early events of HCC development along with feasible time line to study the disease progression. Furthermore, this 6w-DCI model represents a lower hazardous carcinogen treatment regimen (single dose of DEN + 6 weeks of CCl_4_) to induce HCC. It should also be noted that administering CCl_4_ for 4 more weeks (as in the 10w-DCI model), significantly enhances the HCC development compared to age-matched 6w-DCI mice. Thus, comparing various factors between 6w-versus 10w- DCI model might help us gain more insights on the factors driving the disease progression.

To test whether HCC can be induced further rapidly in mice, we sought to increase the dose of DEN. We assessed the effect of DEN dosage on tumorigenesis by injecting increasing dose of DEN (20, 40, and 60 mg/kg) at the ages of 2^nd^, 4^th^, and 6^th^ weeks respectively (41). This is followed by either 6w or 10w of CCl_4_ treatment from the 8^th^ week (Fig. S1C(ii-iii)). Mice were sacrificed 6 weeks post injection and their tumor burden were observed. There was no significant reduction in the tumor development timing in the liver with the increasing dosages of DEN, as compared to the single dose of DEN (Fig.1A). Here also 6w of CCl_4_ after DEN has induced sub-macroscopic nodules (Fig. S1C(ii)) and the group received 10w of CCl_4_ treatment has shown numerous nodules after sacrifice at the age of 24 weeks (Fig. S1C(iii)). Mice treated with vehicle control didn’t show any deformities in liver morphology (Fig. S1C(i)). Mice receiving only three injections of DEN (Fig. S1C(iv)) exhibited few microscopic lesions, indicating that DEN carcinogen alone is not sufficient to induce HCC rapidly. The exposure to chronic liver damaging agents such as CCl_4_ is necessary for robust development of HCC. Throughout the course of treatment, the mice body weight was also monitored and no reduction in the weight was observed (Fig. S2B). Thus, in DEN+CCl_4_ regimen, HCC tumor size and burden showed positive correlation to the CCl_4_ dosage from 6, 10, and 14 weeks of CCl_4_ treatment. However, increasing the dose of DEN did not enhance the HCC tumor formation indicating that initial mutagenic events may not be the rate limiting for the HCC initiation. It is the promotion agent which determines the disease progression (42, 43).

### HCC progression correlates with liver fibrosis and serum-based liver function markers

The differential tumor burden in the 6w- versus 10w-DCI mice provided a longitudinal setting to investigate pro-tumorigenic factors. Histopathological analysis of H&E images demonstrated characteristic features of well to moderate differentiation (as the mice age increased), including thick trabecular architecture, nodular aggregates, increased nuclear-to-cytoplasmic ratio, and prominent nucleoli, confirming chemically induced HCC development in both the cohorts (Fig. 2A(ii-iii)). A few mice also presented with poorly differentiated tumors in the 10w-DCI cohort (in later time points). The H&E staining of the liver tissues from DEN-three injections model also showed moderately differentiated HCC (Fig. S2A). The CCl_4_ treatment is reported to induce fibrosis in the liver via immune infiltration and inflammation (23). Reticulin fibre network is composed primarily of type III collagen and forms a delicate supporting framework within the liver’s space and maintains hepatocyte plate architecture (39). As HCC progresses, there is dissolution of reticulin meshwork once dysplastic nodule progresses to HCC(39). We observed increased damage to reticulin structure and highly thickened fibrillar deposits around tumor nodules indicating the aggressiveness of the HCC (Fig. 2A(ii-iii)). There was no such changes got reported in control group liver (Fig. 2A(i)). In both the carcinogen cohorts, the reticulin fibre network was completely lost inside the tumor nodules suggesting the unstructured growth of HCC cells. Further, analysis of fibrosis in liver by Picrosirius red (PSR) staining revealed deposition of peritumoral bright red birefringent fibres in treated group indicating mature collagen accumulation (Fig. 2B (ii, iii)), whereas the control group showed no such observations (Fig. 2B(i)). In 6w-DCI mice, collagen accumulation progressively increased with age, initially appearing as fibrillar thickening near ducts and sinusoids and later expanding around tumour margins and throughout the liver. However, fibrosis in 6w-DCI model remained lower compared with 10w-DCI mice which exhibited pronounced fibrotic deposition from earlier stages (Fig. 2C). Furthermore, the process of fibrosis continues even in the absence of promotion agent (CCl_4_) suggesting that tumor growth might support the fibrosis.

**Fig. 2.**
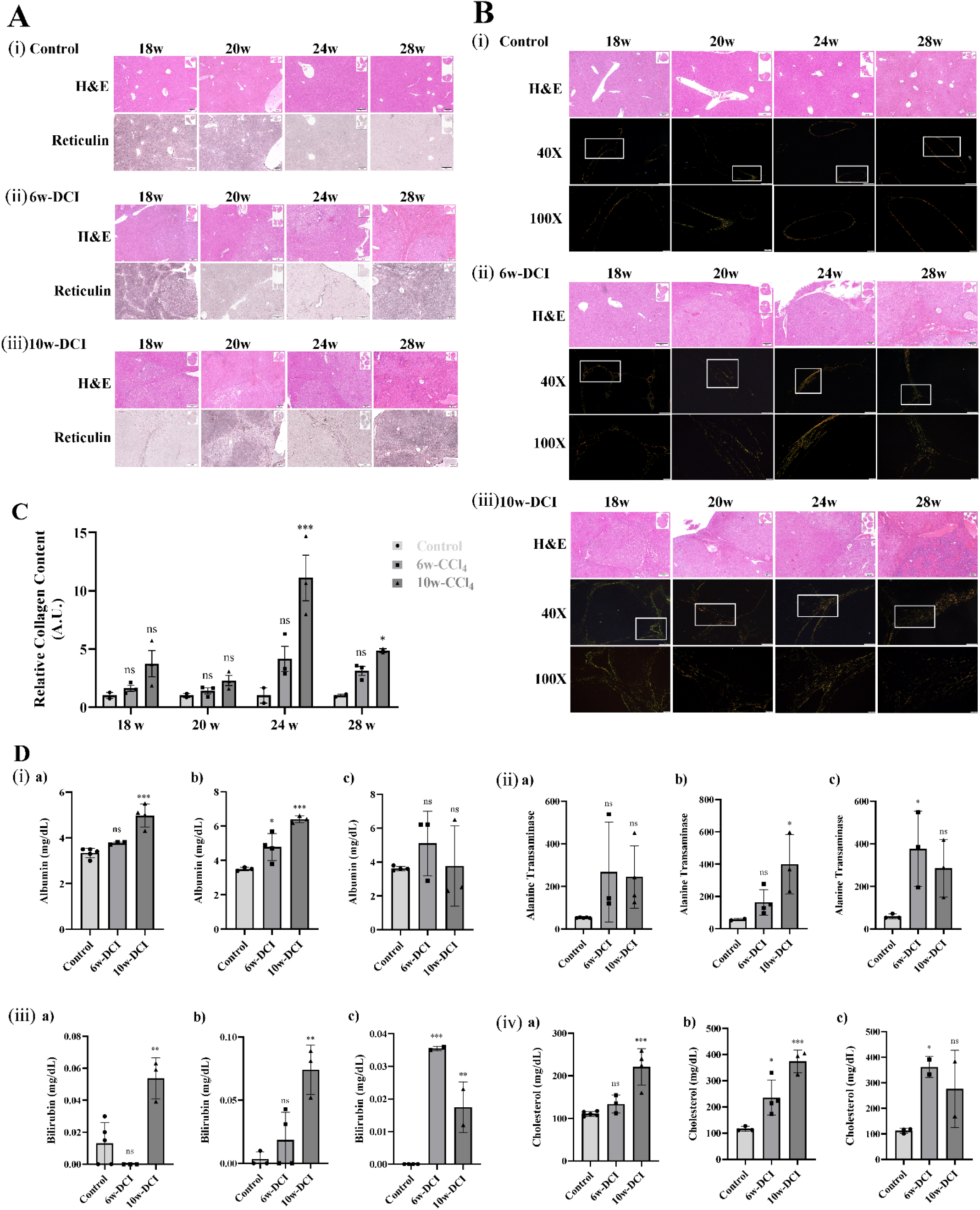
HCC progression is associated with liver fibrosis and serum markers. **A**) Representative reticulin staining images of liver section of tumor shows disruption of reticulin network, (i), Vehicle control; (ii) 6w-DCI; (iii), 10w-DCI; (Scale Bar = 200um). **B**) Picrosirius red staining (PSR) showing progression in collagen deposition during the HCC development as the tumor growth progresses (i) Vehicle control; (ii) 6w-DCI; (iii) 10w-DCI (Scale Bar = 200um) (n=4-6 per time point). A zoomed-in image (100X) of selected region is shown below the 40X image, and H&E staining of same sections are shown above each image. The mice age (in weeks) is mentioned above the H&E sections. **C**) Quantification of the PSR staining. The X-axis shows mouse age. Each dot represents quantification of individual mouse tissue. The values are normalized against control mice of each age group. The *p-values* are calculated from two-way ANOVA tests and, significance marks are indicated above respective bar in comparison to control. **D**) Liver Function assessment in serum. The serum markers were analyzed from control, 6w-DCI, and 10w-DCI mice at **a**)18 weeks, **b**) 24 weeks, and **c**) 28 weeks of age. The biochemical analyses were performed for quantification of (i) Albumin; (ii) Alanine Transaminase; (iii) Bilirubin; and (iv) Cholesterol levels in serum. The Y-axis shows absolute values in each group (n=2-4 per time point/group). The data is plotted as bar-graph and shown as means ± SEM. The *p-values* are from one-way ANOVA tests and, significance marks are indicated above respective bar in comparison to control. *p*-values are denoted as *ns p*>0.05; \**p* ≤ 0.05; \*\**p* < 0.01; \*\*\**p* ≤ 0.001.

The aforementioned longitudinal analysis in DCI model suggest association of fibrosis with tumor burden, however, fail to establish fibrosis as the cause of HCC tumor progression as we did not capture a stage where only significant fibrosis occurred without the tumor formation. The early tumor development showed no significant accumulation of fibrosis. Also, the fibrotic regions were in close proximity to tumor nodules indicating that the cancer cells may be supporting the development of fibrosis. In addition, increasing the dosage of CCl_4_ from 6 weeks to 10 weeks drastically increased the tumor formation as well as the fibrosis. Thus, these findings indicate that prolonged CCl₄ exposure, rather than short-term treatment, drives extensive fibrotic remodelling and liver damage during tumour progression. Further investigations are required to untangle the two events.

As DEN and CCl_4_ are known to induce liver damage, we tested whether HCC progression correlates with the serum markers for liver function tests. The blood serum was collected from the treated and untreated mice immediately after the sacrifice at 18^th^, 24^th^, and 28^th^ weeks of age and the Liver Function Test (LFT) was performed (Fig. 2D). At the early age (18w), serum markers like Albumin, Bilirubin, Cholesterol and Alanine Transaminase levels showed significant increase only in the 10w-CCl₄ cohort while 6w-DCI had no significant increase compared to the control mice (Fig. 2D (i-iv)). At the age of 24^th^ week, 6w-DCI mice also showed increased levels of albumin, alanine transaminase and, cholesterol suggesting that these markers are associated with the tumor growth (Fig. 2D (i-iv)). The bilirubin levels showed significant accumulation at 28^th^ week of age in 6w-DCI cohort as well as 10w-DCI cohort. At the age of 28 weeks, the levels of total protein, albumin and cholesterol did not show much significant difference suggesting that these factors might not be ideal markers for HCC progression. Serum levels of triglycerides, alkaline phosphatase (ALP), aspartate transaminase (AST), and alanine transaminase (ALT) were also measured; however, the observed patterns were not conclusive (Supp. Fig. S2C(i-iv)). Thus, analysis of standard serum markers of liver function is able to represent intermediate progressive phase of HCC, however, their correlation failed to detect early stage of HCC development.

### MAPK and Wnt Signaling are activated in chemical carcinogen induced HCC

To investigate the oncogenic signaling pathways activated in the DCI model, we performed immunoblot analysis for various pathways. Analysis of MAPK downstream targets; MEK and ERK showed increased phosphorylation of MEK1/2 in tumor tissues of both the 6w-and 10w-DCI cohort compared with liver tissues from control mice (Fig. 3A(i-iii)). It should be noted that the total levels of ERK1/2 were increased in tumor tissues, thus, overall phosphorylated forms of ERK1/2 are high in HCC cells. The phosphorylation of p38 was reduced in tumors compared to control, supporting its reported antitumorigenic role in HCC (Fig. 3A(i&iv)). To determine whether these alterations were tumor-specific or a consequence of carcinogen induced liver damage across liver tissue, tumor and adjacent non-tumor regions were analyzed separately. Analysis of tumor and non-tumor regions of 10w-DCI showed increased phosphorylation of MEK1/2 specifically in tumors (Fig. S3A(i)) at the age of 24 weeks. Analysis of β-catenin levels showed increased accumulation in tumor tissues of 6w-DCI group relative to controls (Fig. 3B(i-iii)) indicating the activation of the Wnt/β-catenin pathway. Furthermore, increased phosphorylation of GSK3β at serine 9 was found in the tumors from the 6w-DCI cohort at the age of 24^th^ week (Fig. S3A (ii)). Analysis of tumor and non-tumor regions of 6w-DCI show increased accumulation of β-catenin and phosphorylated-GSK3β in tumors (Fig. S3A(ii)). The activation of Wnt/β-catenin pathway was more prominent in the HCC tumors derived from 3 doses of DEN + CCl_4_ cohort at the 24^th^ week of age (Fig. S3B(i)) along with DEN+CCl_4_ (14w) model (Fig. S3B(ii)). Analysis of tumor and adjacent non-tumor regions obtained from HCC patients reveal the increased phosphorylation of ERK and MEK (Fig. 3C(i)) and activation of Wnt/β-catenin pathway in few patients only (Fig. 3C(ii)). These results suggest that not all patients possess hyperactivation of MAPK and Wnt/β-catenin signaling in HCC.

**Fig. 3.**
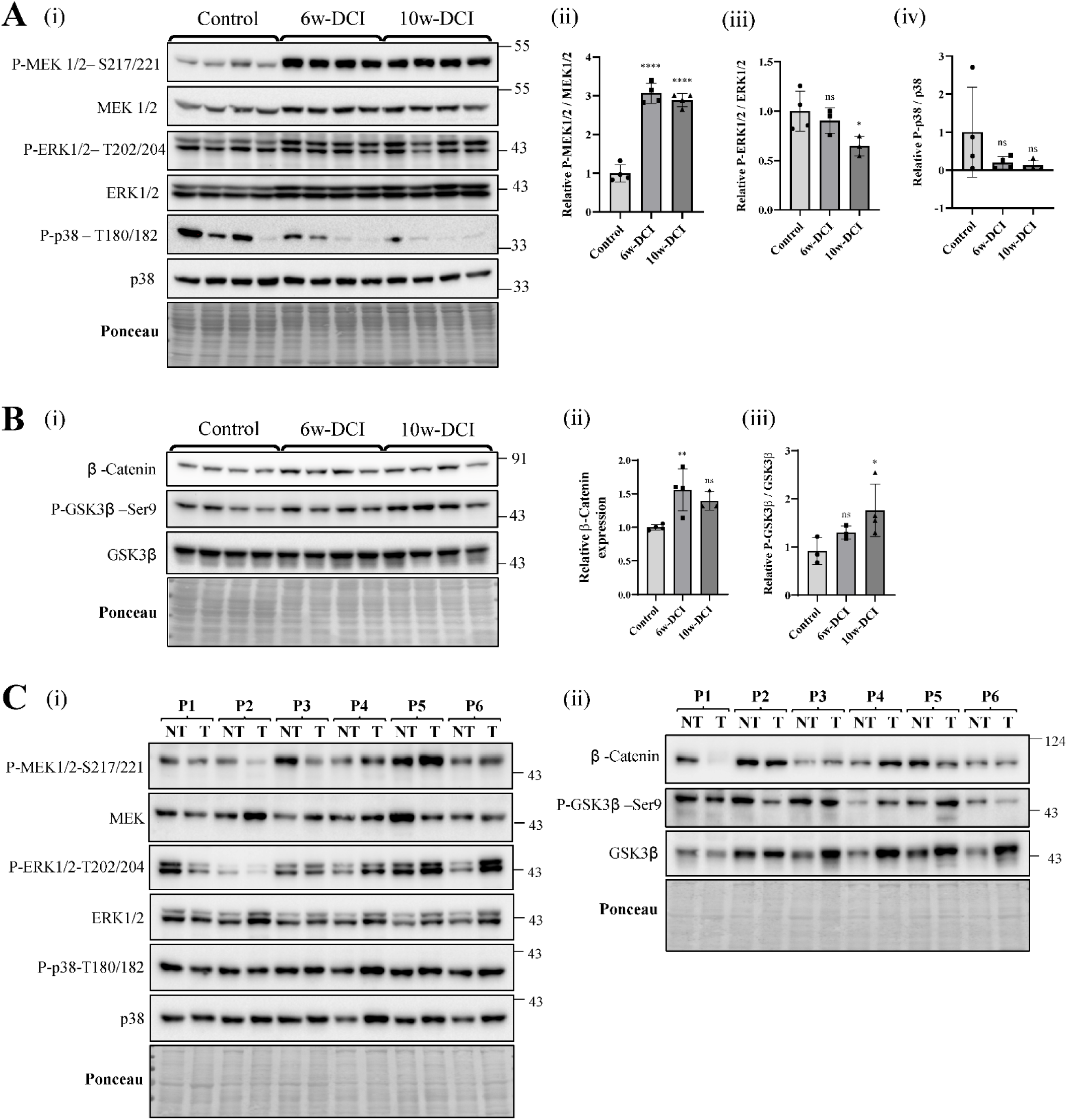
Activation of ERK-MAPK pathway and Wnt/β-Catenin signaling in DCI-HCC mice and HCC patients. **A**) (i) Immunoblot of total and phosphorylated-MEK, ERK, and p38 showing the activity of MAPK pathway in DCI-HCC model. The tissues from 24 weeks old mice belonging to Vehicle, 6w-DCI, and 10w-DCI are compared (n=4 each). The quantification of relative phosphorylation status of P-MEK1/2 (ii), P-ERK 1/2 (iii), and P-p38 (iv) are shown as bar-graph. **B**) (i) Activity estimation of the Wnt/β-Catenin signaling in HCC bearing mice compared to vehicle group. The liver tissues from 24 weeks old mice belonging to Vehicle, 6w-DCI, and 10w-DCI are compared (n=4 each). The quantification of the blots β-Catenin in comparison to total protein (ii) and P-GSK3-β with respect to its total form (iii) are shown as bar-graph. **C**) P-MEK1/2, P-ERK1/2, and P-p38 status and their total levels (i) and Wnt-β Catenin and P-GSK3-β levels (ii) in paired tumor and non-tumor tissues obtained from HCC patients were analyzed by immunoblotting (n=6). In bar plots, each dot represents values from an individual mouse. The data is shown as means ± SEM. The *p-values* are from one-way ANOVA tests and, significance marks are indicated above respective bar in comparison to control. *p*-values are denoted as *ns p*>0.05; \**p* ≤ 0.05; \*\**p* < 0.01; \*\*\*\**p* < 0.0001.

### Deregulation of mTOR pathway in HCC

The PI3K-AKT-mTOR axis is upregulated in majority of HCC cases (44). The immunoblot analysis of mTORC1 downstream target S6K showed reduced phosphorylation at T389 residue (Fig. 4A) in HCC tumors from both 6w- and 10w-DCI cohorts (at 24 weeks age). Accordingly, the phosphorylation of RPS6, the downstream target of S6K is also reduced in these tumors (Fig. 4A) indicating suppression of mTORC1-S6K-S6 axis in DCI-HCC. Immunoblot quantification revealed a reduction of 4-fold in S6K activity followed by nearly 2-fold in S6 and AKT phosphorylation compared to control livers (Fig. S4C(i-iii)). The analysis of tumor and non-tumor region of mice liver tissues also confirmed the tumor-specific downregulation of mTORC1-S6-axis (Fig. S4A(i)). Analysis of tumor and non-tumor region of HCC patients also revealed reduced phosphorylation of RPS6 in tumor-specific manner (Fig. 4B). The autophosphorylation of mTOR at Serines 2440/2448 suggested no reduction indicating that the activity of mTOR may not have directly impacted (Fig. S4A(i)). The mTORC2-mediated phosphorylation of AKT at S473 is slightly reduced in the DCI-derived HCC tumors (Fig. 4A). However, in patient-derived HCC tissues did not show downregulation of AKT phosphorylation at 473 (Fig. 4B). Thus, these results reveal unanticipated observation of reduced mTORC1-S6K-S6 axis in HCC tumors.

**Fig. 4.**
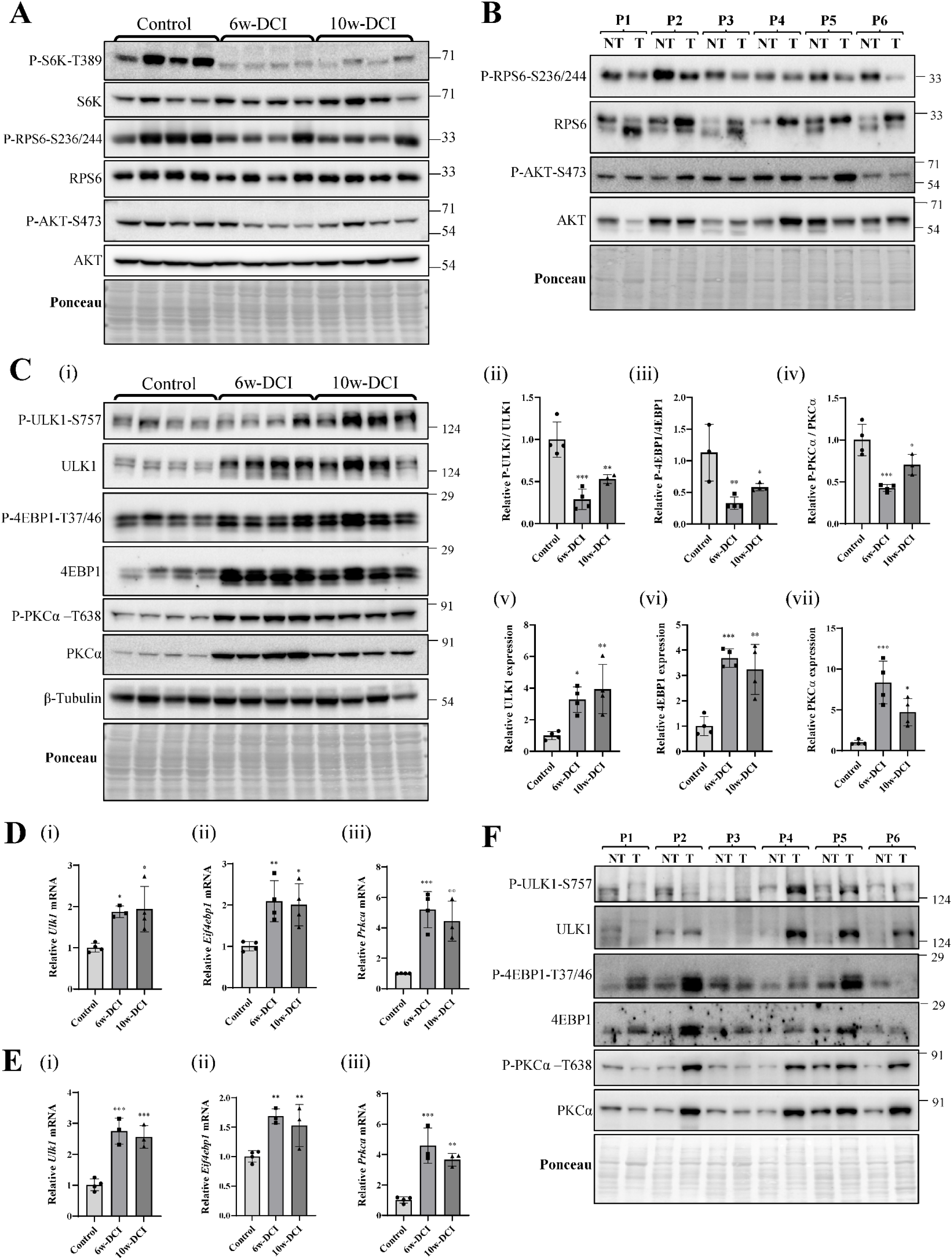
Deregulation of mTOR signaling in DCI-HCC mice and HCC patients. **A**) Immunoblots of P-S6K/S6K, P-RPS6/RPS6 and P-AKT/AKT in liver tissues obtained from vehicle, 6w-DCI, and 10w-DCI mice sacrificed at the age of 24 weeks (n=4 each). **B**) The immunoblot analysis of P-RPS6/RPS6 and P-AKT/AKT in the paired tumor and non-tumor tissues from HCC patients (n=6). **C**) (i) Immunoblotting of P-ULK1/ULK1, P-4EBP1/4EBP1, and P-PKCα/PKCα in liver tissues obtained from vehicle, 6w-DCI, and 10w-DCI mice sacrificed at the age of 24 weeks (n=4 each). The relative quantification of P-ULK1/ULK1 (ii), P-4EBP1/4EBP1 (iii), and P-PKCα/PKCα (iv) are shown as bar diagram. The relative total levels of ULK1 (v), 4EBP1 (vi), and PKCα (vii) were measured with respect to β –tubulin and shown as bar diagram. **D**) The transcript levels of *Ulk1* (encoding ULK1) (i), *Eif4ebp1* (encoding 4EBP1) (ii) and *Prkca* (encoding PKCα) (iii) were measured by quantitative real-time PCR (qRT-PCR) in control, 6w-DCI, and 10w-DCI mice sacrificed at the age of 24 weeks (n=3-4 each). The transcript levels were normalized to *Gapdh* levels and the relative quantification with respect to vehicle control are shown. **E**) The transcript levels of *Ulk1* (i), *Eif4ebp1* (ii) and *Prkca* (iii) are measured by quantitative real-time PCR (qRT-PCR) in control, 6w-DCI, and 10w-DCI mice sacrificed at the age of 28 weeks (n=3-4 each). Data is plotted as bar-graph representation of values. Data is represented as means ± SEM. *p-values* are from the one-way ANOVA test and significance marks are indicated above respective bar in comparison to control. **F**) Expression analysis of P-ULK1/ULK1, P-4EBP1/4EBP1 and P-PKCα/PKCα in HCC in the paired tumor and non-tumor tissues from HCC patients (n=6). *p*-values are denoted as \**p* ≤ 0.05; \*\**p* < 0.01; \*\*\**p* ≤ 0.001.

As mTOR signaling has several downstream mediators, we investigated the phosphorylation status of other major regulators. The analysis of mTORC1 targets 4EBP1, and ULK1; and mTORC2 target PKCα showed significant increase in the phosphorylation in tumor tissues relative to control liver (Fig. 4C(i)). Notably, the total protein level of ULK1 and 4EBP1 is increased to 3-4-fold, and PKCα is increased >5 fold in both 6w- and 10w-DCI models at the 24 weeks of age (Fig. 4C(v-vii)). The qPCR analysis showed nearly 2-fold upregulation of *Eif4ebp1* (encoding 4EBP1) and *Ulk1* (encoding ULK1) transcripts, and around 4-fold upregulation of *Prkca* (encoding PKCα) transcripts in tumor tissues compared with normal liver at both 24 weeks (Fig. 4D(i-iii)) and 28 weeks (Fig. 4E(i-iii)) of age, indicating that the mechanism of upregulation of these substrates to be transcriptional. The total levels of 4EBP1, ULK1, and PKCα are elevated in the 14w-DCI model (Fig. S4B (i)) along with DEN-three DCI model (Fig. S4B (ii)). The analysis of tumor and non-tumor regions showed tumor-specific upregulation of 4EBP1, ULK1, and PKCα in the DCI model (Fig. S4A (ii)).

Analysis of patient-derived HCC tissues and their adjacent non-tumor region also showed similar upregulation of total levels of ULK1, 4EBP1 and PKCα in tumor regions (Fig. 4F). To obtain the effect of ULK1, 4EBP1 and PKCα on HCC patient survival, Kaplan–Meier survival analysis was performed using TCGA liver hepatocellular carcinoma (HCC; n=245) and pan-cancer cohorts (n=4639) stratified by ULK1, EIF4EBP1 and PRKCA expression. The median follow-up duration for the cohort was 19.58 months (ranges from 0.03 to 120.7 months) and during this period, 179 Recurrence and 130 fatal events occurred. In HCC, elevated EIF4EBP1 was associated with a rapid decline in 5-year progression-free survival (PFS) (Fig. S4D(i)), consistent with its role in disease aggressiveness, and similarly showed a broad oncogenic association in pan-cancer analysis (Supp. Fig. 5A(i)). ULK1 upregulation demonstrated a strong association with poor PFS in HCC (Fig. S4D(ii)), but only a modest effect across cancers (Supp. Fig. 5A(ii)), suggesting its crucial role in the development of liver tumors. PRKCA (PKCα) overexpression showed a robust and significant association with poor PFS in both HCC and pan-cancer cohorts (Fig. S4D(iii); Fig. S5A(iii)). Median 5-year overall survival suggested a potential role for ULK1 in disease progression; however, this was not statistically significant, likely due to limited sample size and follow-up constraints (Fig. S5B (ii)) The other two molecules haven’t shown much correlation with the 5-year overall survival (Fig. S5B (i, iii)). Also, the median hazard ratio of EIF4EBP1 and ULK1 expression was also found to be more in case of HCC patients as compared to the overall group (Fig. S5C)

**Fig. 5.**
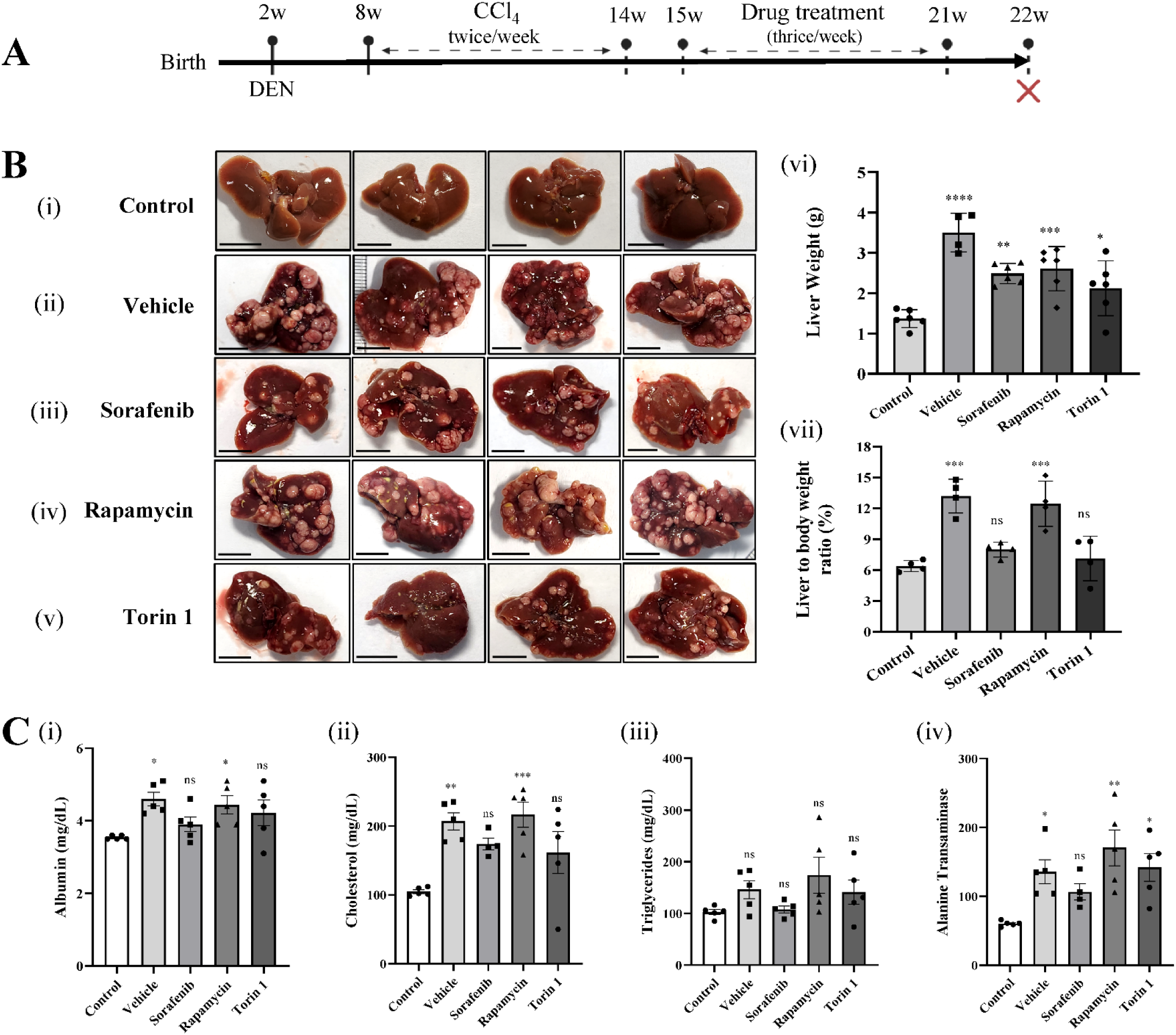
Inhibition of both mTOR complexes reduces HCC progression. **A**) Schematics of HCC induction and drug treatment regimen in DCI model. The 6w-DCI mice were treated with sorafenib, Rapamycin and Torin 1, vehicle group received placebo (DMSO). (n=6 each). **B**) (i – v) Representative liver images of different groups. (Scale bar = 1cm), (vi) Liver weight quantification at sacrifice for mice of different groups (n=6 each) along with (vii) Liver to Body weight ratio of mice (in percentage) received different treatments (n=4 each). Data is represented as means ± SEM and plotted as bar graph representation of weight values (g). **C**) Blood serum profile showing liver functioning in HCC bearing mice treated with different drugs, (i) Albumin; (ii) Cholesterol levels; (iii) Triglycerides; and (iv) Alanine Transaminase, (n=3-5 per group), Data is represented as means ± SEM and plotted as bar graph representation of absolute values. Each dot represents individual mouse. The *p-values* are from one-way ANOVA tests and indicated above respective bar in comparison to control. *p*-values are denoted as *ns p*>0.05; *\*p* ≤ 0.05; *\*\*p* < 0.01; *\*\*\*p* ≤ 0.001; *\*\*\*\*p* < 0.0001.

### Blocking both mTOR complexes reduces HCC progression

As the above-mentioned data indicate that HCC possesses deregulated biased mTOR signaling, we further investigated whether mTOR signaling plays any role in HCC progression in the DCI model. To test the impact of mTOR pathway in the HCC progression, 6w-DCI mice were treated with mTORC1 inhibitor Rapamycin or mTORC1 and mTORC2 dual inhibitor Torin 1 (Fig. 5A). The 6w-DCI mice at were treated with 10mg/kg of each drug from 15^th^ week, thrice per week for 6 weeks and the mice were sacrificed post 1 week of treatment regimen (at the age of 22^nd^ week). Compared to the vehicle group (Fig. 5B(ii)), Rapamycin treatment was not able to inhibit HCC tumor growth (Fig. 5B(iv)). However, the Torin 1 treatment resulted in lesser tumor burden (Fig. 5B(v)) in comparison to the vehicle. Sorafenib is an FDA approved multikinase inhibitor for HCC and it significantly reduced the HCC progression (Fig. 5B(iii)). The measurement of liver weight and the ratio of liver to body weight shows significant reduction upon Torin 1 or Sorafenib treatment (2mg/kg) while Rapamycin did not (Fig. 5B(vi-vii)); indicating that the mTOR dual kinase inhibitor is effective in blunting HCC progression. The control cohort indicates the healthy mice without induction of HCC (Fig. 5B(i)). The serum biomarker analyses showed a reduction in the albumin, cholesterol, triglycerides, and alanine transaminase levels upon treatment with Sorafenib and Torin 1 treatment as they were comparable to control mice while vehicle treated 6w-DCI mice showed significant elevation of these markers (Fig. 5C(i-iv)). The other serum markers such as total protein and aspartate transaminase did not show any meaningful pattern between the treated and untreated 6w-DCI mice at the age of 22^nd^ week (Fig. S5D(i-ii)). The rapamycin treatment alone did not reduce these serum markers suggesting that inhibition of mTORC1 alone may not be sufficient to inhibit HCC progression. Thus, these results indicate that targeting both mTORC1 and mTORC2 complexes may have better therapeutic potential against HCC.

## Discussion

The upregulation of mTOR signaling in HCC mainly occurs via loss-of-function mutations in TSC or PTEN or via hyperactive mutations in PI3K (9, 36, 45). Although these genetic alterations are rare, phosphoproteomic studies suggest hyperactivation of mTOR in 40-50% HCC cases (44). However, our DCI model as well as the HCC patient tissue analysis reveal decreased activity of mTORC1-S6K-S6 axis. Interestingly, tumour tissues displayed elevated expression and phosphorylation of key downstream effectors, including 4EBP1, ULK1, and PKCα. The tumour-specific upregulation of 4EBP1, ULK1, and PKCα, observed in both mouse and human HCC samples, underscores their potential as critical mediators of tumour adaptation under stress conditions (46–48). Importantly, elevated expression of these molecules correlated with poor progression-free survival in HCC patients, supporting their clinical relevance. Role of upregulated expression of 4EBP1(49–51) and ULK1(52–54) in tumor growth and metastasis is observed in multiple cancer types, suggesting their clinical significance and importance as druggable targets. This paradox suggests the presence of compensatory or alternative regulatory mechanisms that sustain selective mTOR signaling branch to support oncogenic demands.

mTOR signaling, especially the mTORC1 is the homeostatic signaling pathway which balances cell growth and physiological functions under favourable growth conditions and the energy supply (34). For cancer cells, this may become burden as not all the cellular jobs have to be taken care by a cancer cell. For example, cancer cells may take up most of the metabolites instead of synthesizing them (55, 56). Thus, skewing mTOR activity towards specific branches may provide an efficient mechanism to cope with the hostile environment of cancer progression. Thus, elicitation of biased mTOR signaling may serve as an oncogenic mechanism for HCC cells. The biased signaling has been reported in the context of GPCR signaling (57). Our study reports the substrate abundance-mediated biased mTOR signaling in HCC. Similar biased signaling may occur in other cancers and disease contexts which may be of future interest. The molecular mechanisms of elicitation and the oncogenic role of biased mTOR signaling demands more future investigations. Recent studies suggest spatial compartmentalization-mediated mTOR signaling and whether this contributes for the observed biasness will be interesting to explore (58, 59).

The pre-clinical animal models are crucial for the understanding of cancer initiation, progression and assessment of novel therapeutic regimens. However, the long duration for developing such models daunt the molecular investigations. The chemical-induced carcinogen models of HCC are instrumental in testing various therapeutic strategies. However, their use in basic understanding of disease progression is limited due to heterogeneity in the oncogenic drivers and lack of proper longitudinal models. Our study establishes a relatively low invasive strategy using DEN + CCl_4_ to develop HCC in mice (6w-DCI model) by single dose of DEN at 2^nd^ week and 6 weeks of CCl_4_ treatment from 8^th^ week of age. This model develops HCC within 4 months and can be useful to study early events of HCC tumor formation and monitor their progression. The prior reports have developed HCC using DEN + CCl_4_ and have shown to be ideal for fibrosis and liver damage induced HCC (23, 25). Our serum marker analysis also showed accumulation of cholesterol and triglycerides upon development of HCC, indicating the metabolic dysregulation. The advantage of 6w-DCI model is that the HCC development occurs post carcinogen treatment regimen, thus, drastically reducing the confounding impact of carcinogen itself. This model can also be useful in studying effect of various drugs which can combat HCC progression as Sorafenib, an FDA approved drug could inhibit the HCC progression (Fig. 5B(iii)).

Apart from revealing biased mTOR signaling, the DCI-model recapitulates key oncogenic signaling alterations observed in human HCC, including activation of MAPK and Wnt/β-catenin pathways. The genomic analysis of DEN-induced HCC has shown heterogeneous mutations in BRAF and APC to induce these pathways(2). In agreement, our model also shows increased MEK phosphorylation and β-catenin accumulation, indicating the activation of proliferative and survival pathways. Interestingly, the MEK-mediated phosphorylation of ERK is reduced relative to total ERK levels. Whether hyperphosphorylated MEK directly exerts its oncogenic transcriptional programs or it has any other protein substrates is interesting to explore. The phosphorylated p38 is known to induce apoptosis and cell death, thus, acting as tumor suppressor. Reduction in p38 phosphorylation supports the tumor growth.

Our study reveals that both mTORC1 and mTORC2 play crucial role in progression of HCC. Blocking only mTORC1 with Rapamycin does not lower tumour burden, but blocking both mTORC1 and mTORC2 with Torin 1 greatly reduces tumour growth, similar to the effects of sorafenib. The effect of rapamycin in terms of differential inhibition of mTORC1 substrates such as S6K and 4EBP1 is previously established due to the differential sterical hindrance caused by rapamycin-FKBP12 complex (60, 61). Rapamycin is well known as immune suppressor and thus, in case of carcinogen induced HCC development, maybe use of Rapamycin reduces the war between the cancer cell growth versus immune cells. Furthermore, recent studies implicate important role of mTORC2 in oncogenic lipogenesis and cancer cell metastasis(62). In addition, mTORC2 regulates glucose metabolism and ROS metabolism which in turn drive tumor cell survival (63). Thus, blocking both mTOR complexes with inhibitors such as Torin 1 may have better therapeutic potential. Future research will help in further understanding the role of biased mTOR signaling and better combination therapies instead of targeting the master regulator of cell growth and metabolism.

## Abbreviations

HCC: Hepatocellular carcinoma
DEN: diethyl nitrosamine
CCl_4_: carbon tetrachloride
DCI: DEN-CCl_4_ induced

## Acknowledgements

The authors thank all the Shetty lab members for their input on the manuscript. RS is supported by a fellowship from the Department of Biotechnology (DBT)-Government of India. NS is supported by ANRF National Postdoctoral fellowship. PM and AS are supported by fellowships from the ACTREC, Tata Memorial Centre. This research is supported by DAE DPR sanction number 1/3(4)/2021/TMC/R&D-II/15063, ICMR intermediate grant IIRPG-2023-0137 and ICMR small grant IIRPSG-2025-01/04589/SG. We thank Mr. Ganesh Dahimbekar and Dr. Amartya Singh for analysis of TCGA dataset. We also thank composite laboratory, pathology, surgical pathology, and Laboratory animal facility of ACTREC for all the support. The schematics were drawn using BioRender.

---

**Fig. S1.**
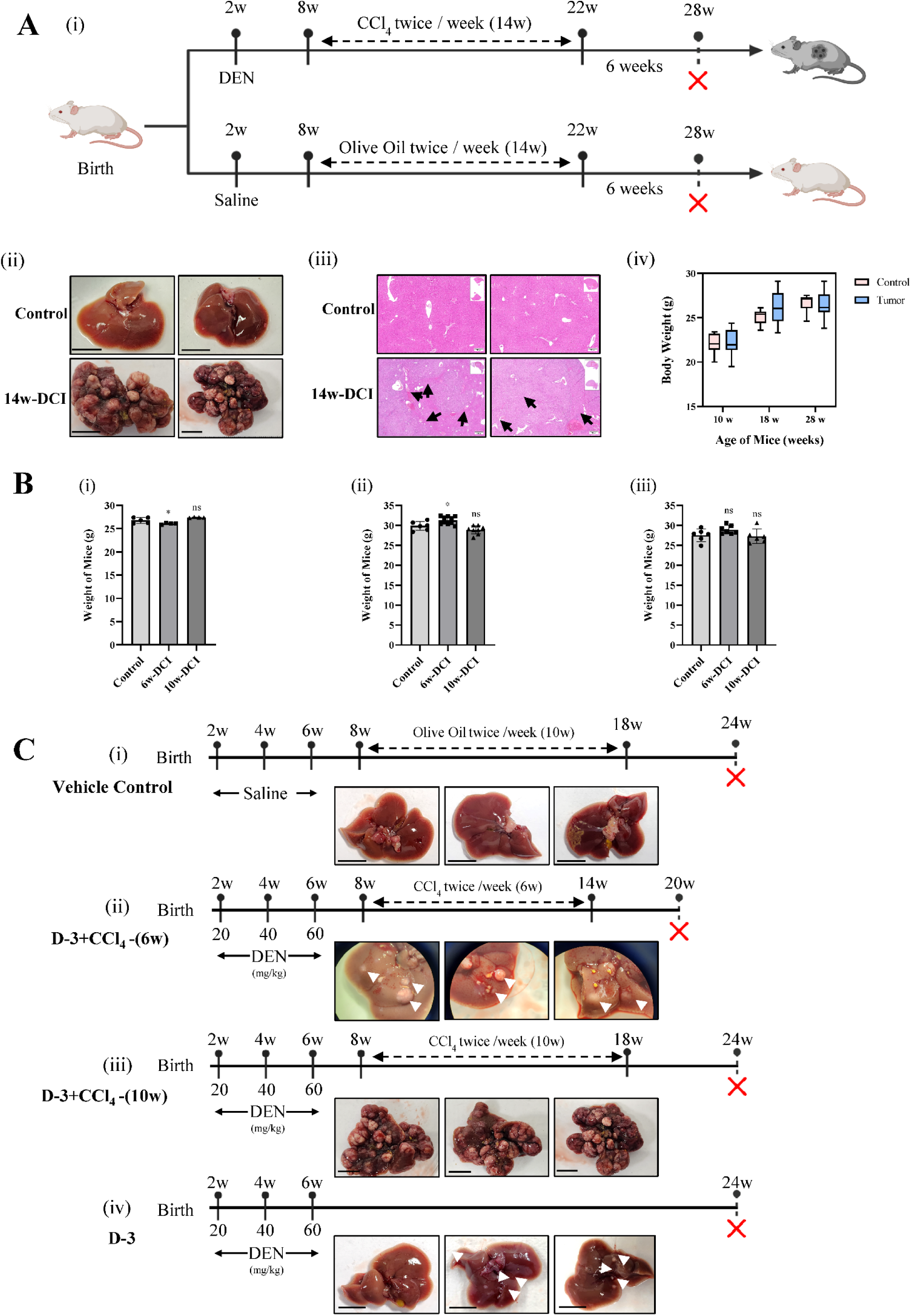
Enhanced genotoxicity via DEN has no effect on rapid tumorigenesis. **A**) (i) Schematics of HCC tumor establishment, referred as 14w-DCI. (ii) Representative liver images of mice from control group and 14w-DCI group (n=10-12) (Scale Bar = 1cm), followed by their H&E imaging (Scale Bar = 500µm) (iii), tumors marked with arrows. (iv) Body weight quantification of mice at different time points. **B**) Body weight estimation of 6w-DCI and 10w-DCI mice at 18^th^, 24^th^, and 28^th^, weeks of age against vehicle control group. Data is represented as means ± SEM and plotted as bar graph representation of weight values (g). Each dot represents liver weight of single mouse. The *p-values* are from one-way ANOVA tests and indicated above respective bars in comparison to control. **C**) Carcinogen treatment schema in mice for the increased DEN dosage cohort followed by their representative liver images (Scale Bar = 1cm) (i) Vehicle Control, (ii) DEN 3 dosage + 6w-CCl_4_ treatment group, (iii) DEN 3 dosage + 10w-CCl_4_ treatment group (iv) DEN only treatment, (n=10 each). *p*-values are denoted as *ns p*>0.05; *\*p* ≤ 0.05.

**Fig. S2.**
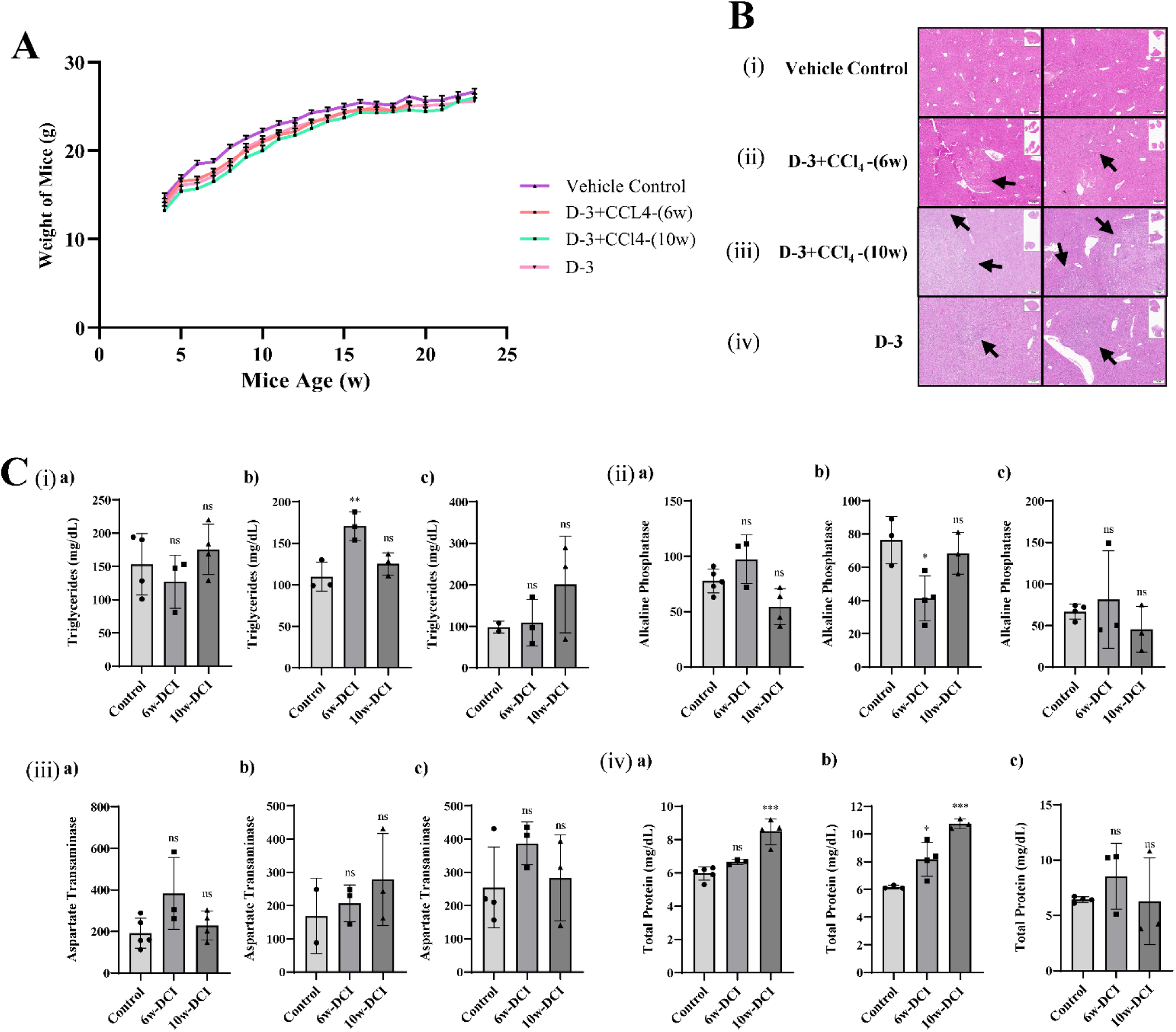
Histological assessment of tumors from increased DEN dosage cohort. **A**) H&E images of liver sections of mice belonging to the increased DEN dosage cohorts with variable treatments, tumor regions are marked with arrows (Scale Bar=200µm). **B**) Body weight quantification of the mice throughout the course of study (n=10 each). **C**) Serum biomarkers assessment of 6w-DCI and 10w-DCI mouse against vehicle control group at different time points, a) 18w, b) 24w, and c) 28w respectively, graphs show (i) Triglycerides; (ii) Alkaline Phosphatase; (iii) Aspartate transaminase; and (iv) Total Protein levels. Data is plotted as bar graph representation of absolute values. Each dot represents a value from single mouse. The *p-values* are from one-way ANOVA tests and indicated above the respective bars in comparison to control. *p*-values are denoted as *ns p*>0.05; \**p* ≤ 0.05; \*\**p* < 0.01; \*\*\**p* ≤ 0.001.

**Fig. S3.**
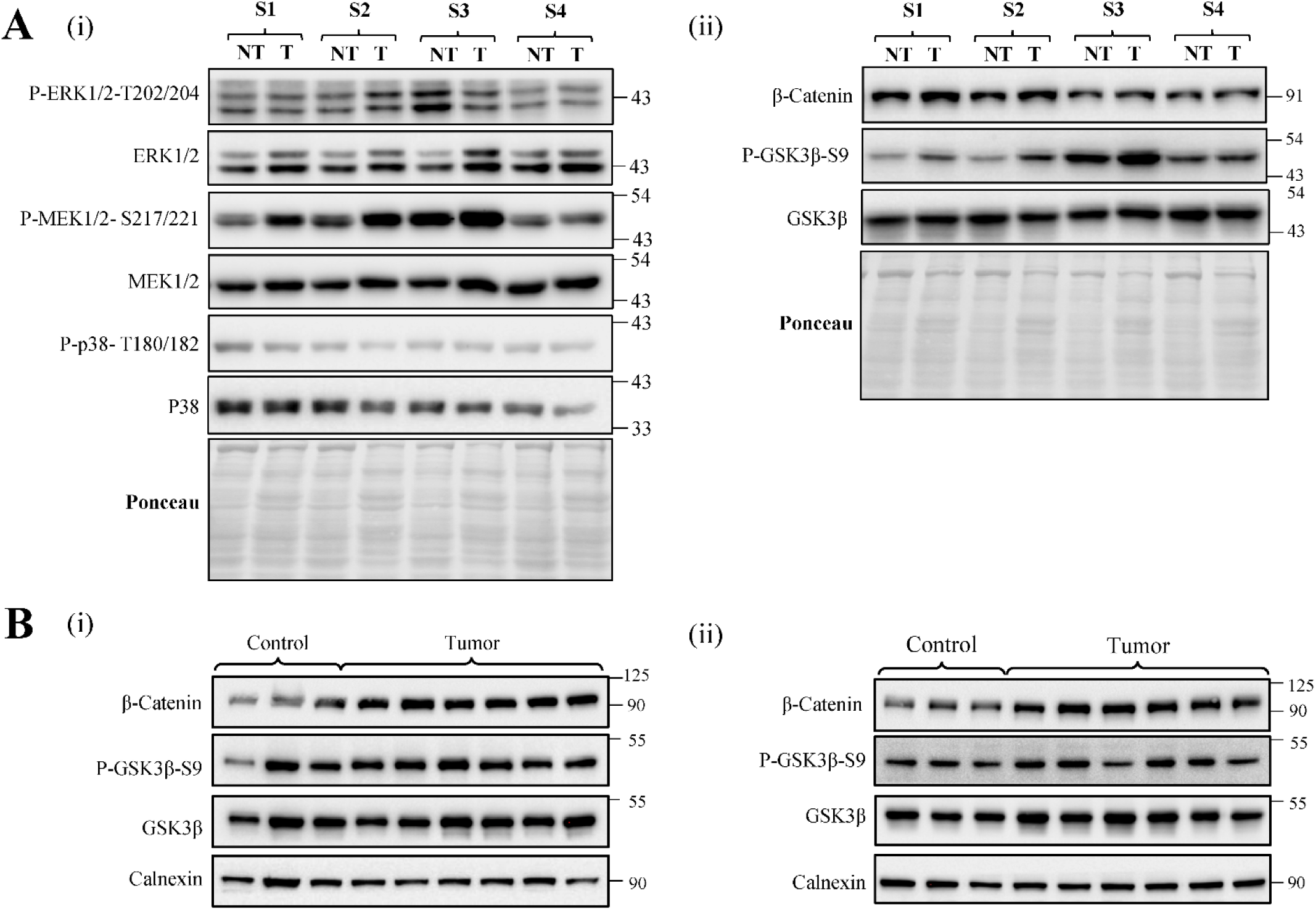
Tumor-specific activation of MAPK and Wnt signaling pathways in 6w-DCI and 10w-DCI group. **A**) (i) Immunoblot of MEK, ERK, and p38 showing the activity of MAPK pathway in tumor and adjacent non-tumor region of same HCC containing livers. (ii) Assessment of Wnt signaling in tumor and adjacent non-tumor region of same HCC containing livers. Samples were named as S1, S2, S3, and S4. Expression is normalized against ponceau staining. **B**) (i) Wnt Signaling analysis in DEN-3 + CCl_4_ (10w) and (ii) 14w-DCI tumors against the control livers. Expression normalization was performed against Calnexin.

**Fig. S4.**
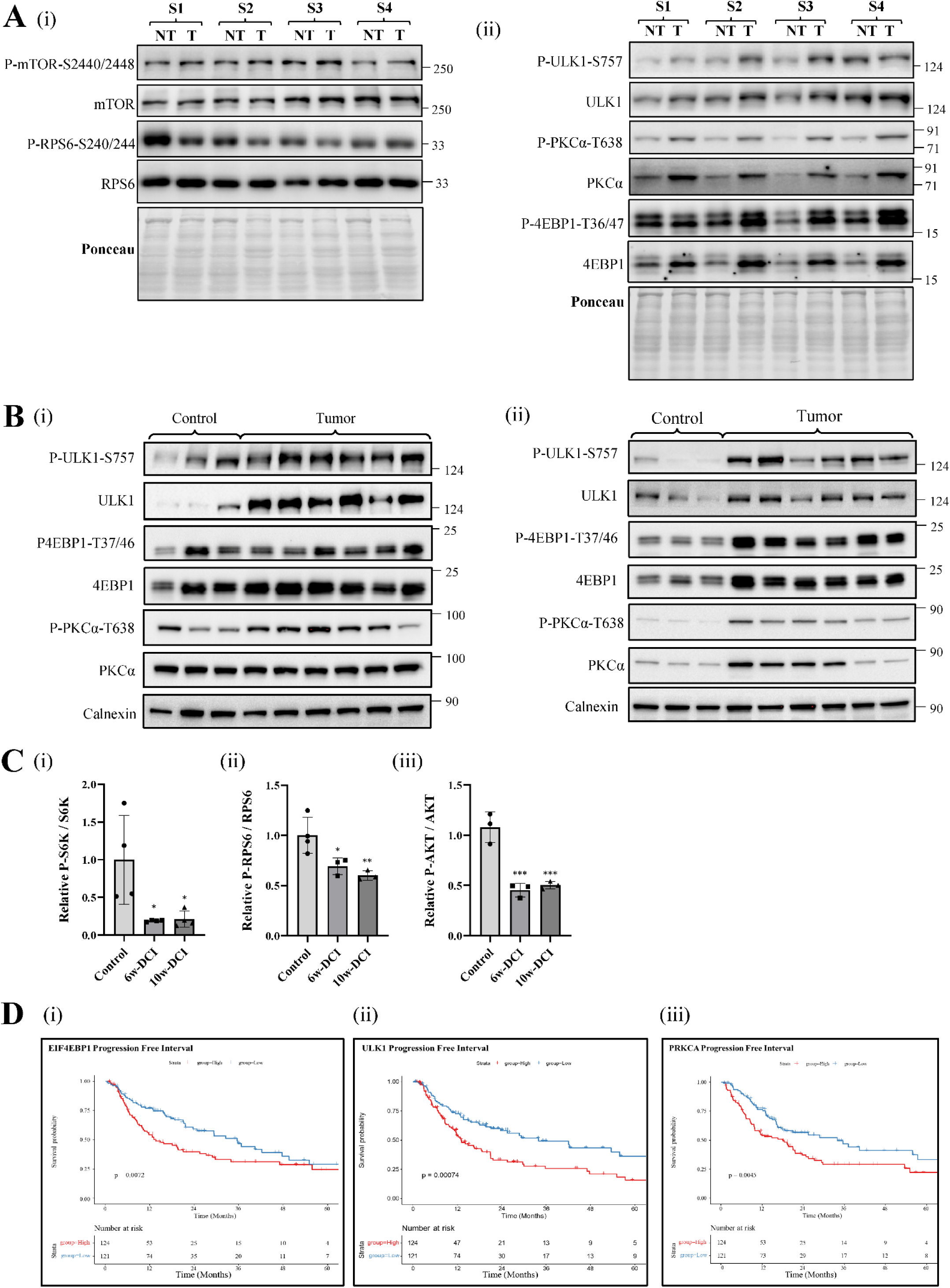
Tumor selective expression of deregulated mTOR pathway substrates in 6w-DCI model correlates with a poor progression free survival in HCC patients. **A**) (i) immunoblotting of P-RPS6/S6 and P-mTORC1/mTORC1, in tumor and adjacent non-tumor region of same HCC containing livers. (ii) Protein based expression changes of P-ULK1/ULK1, P-4EBP1/4EBP1 and P-PKCα/PKCα in in tumor and adjacent non-tumor regions of HCC livers. Samples were named as S1, S2, S3, and S4. Expression is normalized against ponceau staining. **B**) (i) mTOR pathway substrates were analyzed in DEN 3 - CCl_4_(10w) and (ii) 14w-DCI tumors against the control livers. Expression normalization was performed against Calnexin. **C**) (i – iii) Immunoblot quantification of mTOR pathway substrates showing their downregulation in 6w-DCI and 10w-DCI mouse against normal liver. Data is plotted as bar graph representing relative values in tumor group w.r.t. control group. Each dot represents a value from an individual mouse. The *p-values* are from one-way ANOVA tests and indicated above each graph in comparison to control. *p*-values are denoted as *ns p*>0.05; \**p* ≤ 0.05; \*\**p* < 0.01; \*\*\**p* ≤ 0.001. **D**) Median 5-year Progression Free Survival for the HCC Patients expressing either high or low mRNA levels of the markers of the mTOR pathway (i) ULK1 (ii) 4EBP1, and (iii) PKCα, (n=489).

**Fig. S5.**
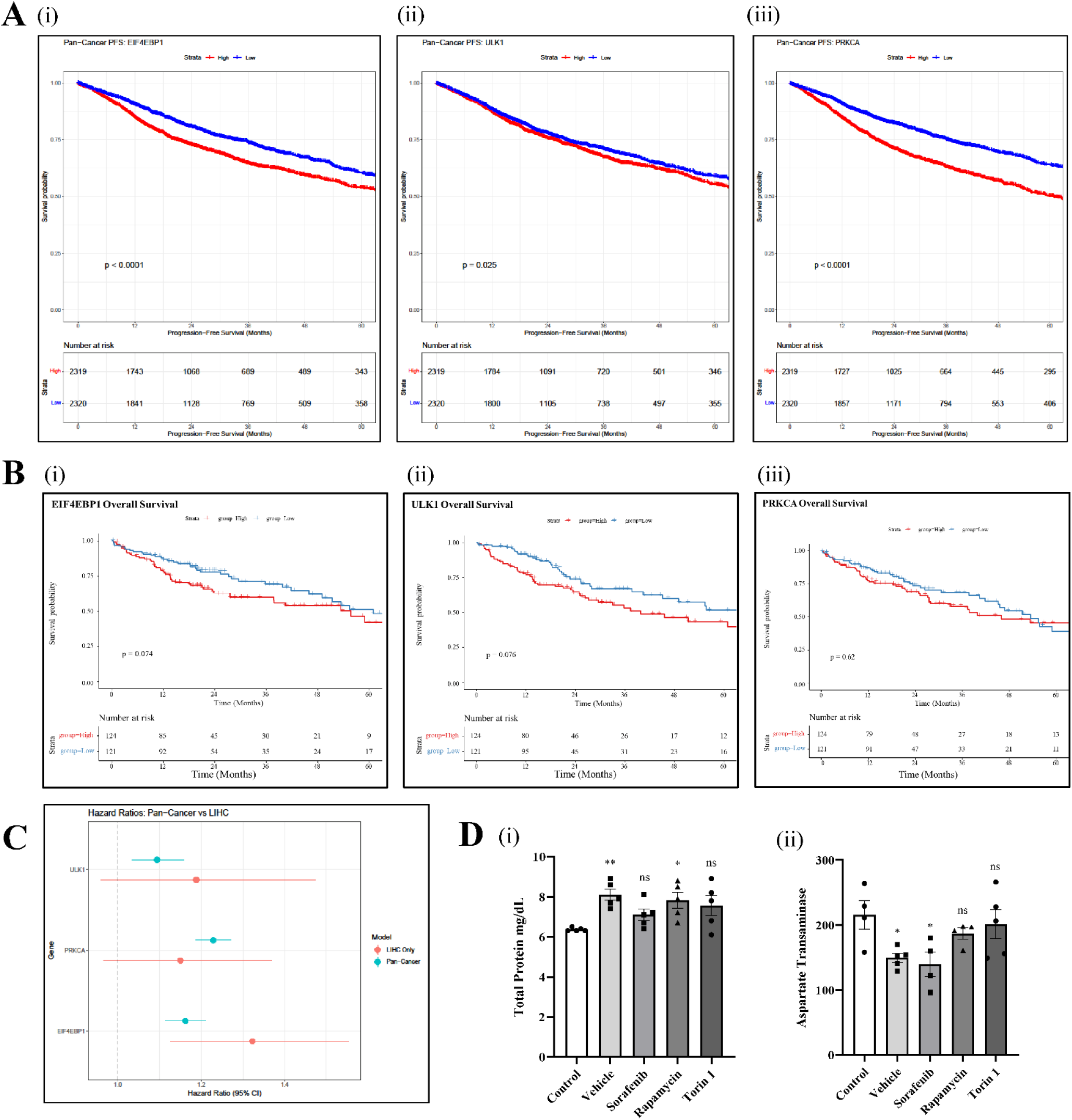
mTOR pathway substrates are tumorigenic in HCC biased condition. **A**) Median 5-year survival for HCC patients according to the mRNA expression of PKCα (i), 4EBP1 (ii), and ULK1 (iii), (n=245). **B**) Impact of expression of mTOR substrates on median 5-year Progression Free Survival of HCC vs Pan-cancer group, (n=4639). **C**) Hazard ratios of PKCα, 4EBP1, and ULK1 in HCC vs Pan-cancer group. **D**) Blood serum profile showing liver function in 6w-DCI-HCC mice treated with various drugs, (i) Alkaline Phosphatase; and (ii) Aspartate transaminase; (n=3-5 per group). Data is plotted as bar graph representing absolute values. Each dot represents a value from an individual mouse. The *p-values* are from one-way ANOVA tests and indicated above each graph in comparison to control. *p*-values are denoted as *ns p*>0.05; \**p* ≤ 0.05; \*\**p* < 0.01.

**Supplementary Table 1.** List of Antibodies used for Immunoblotting.

| Sl. No. | Antigen Name | Company Name | Cat. No. | Dilution. Factor |
| --- | --- | --- | --- | --- |
| 1 | Phospho-p44/42 MAPK (Erk1/2) (Thr202/Tyr204) (E10) | CST | 4696 | 1:1500 |
| 2 | p44/42 MAPK (Erk1/2) | CST | 4695 | 1:2000 |
| 3 | Phospho-MEK1/2 (Ser217/221) (41G9) | CST | 9154 | 1:1500 |
| 4 | MEK 1/2 (D1A5) | CST | 8727 | 1:2000 |
| 5 | Phospho-p38 MAPK (Thr180/Tyr182) | CST | 9211 | 1:1500 |
| 6 | p38 MAPK | CST | 9212 | 1:2000 |
| 7 | Non-phospho (Active) beta-Catenin (Ser33/37/Thr41) | CST | 4270 | 1:3000 |
| 8 | Phospho-GSK-3 beta (Ser9) (D85E12) | CST | 5558 | 1:1500 |
| 9 | GSK-3 beta (D5C5Z) | CST | 12456 | 1:3000 |
| 10 | Phospho-p70 S6 Kinase (Thr389) (108D2) | CST | 9234 | 1:1500 |
| 11 | p70 S6 Kinase | CST | 9202 | 1:2000 |
| 12 | Phospho-S6 Ribosomal Protein (Ser240/244) (D68F8) | CST | 5364 | 1:1500 |
| 13 | S6 Ribosomal Protein (5G10) | CST | 2217 | 1:2000 |
| 14 | Phospho-Akt (Ser473) | CST | 9271 | 1:1000 |
| 15 | Akt (pan) (11E7) | CST | 4685 | 1:3000 |
| 16 | Phospho-ULK1 (Ser757) (D7O6U) | CST | 14202 | 1:1500 |
| 17 | ULK1 (D8H5) | CST | 8054 | 1:2000 |
| 18 | Phospho-4E-BP1 (Thr37/46) (236B4) | CST | 2855 | 1:1500 |
| 19 | 4EBP1 | CST | 9452 | 1:2000 |
| 20 | Phospho-PKC alpha/beta II (Thr638/641) | CST | 9375 | 1:1500 |
| 21 | PKC-alpha | CST | 2056 | 1:2000 |
| 22 | β-Tubulin (D-10) | SCBT | SC-5274 | 1:1000 |
| 23 | Phospho-mTOR (Ser2448) (49F9) | CST | 2976 | 1:1000 |
| 24 | mTOR (7C10) | CST | 2983 | 1:1500 |
| 25 | Calnexin polyclonal antibody | ENZO | ADI-SPA-860 | 1:5000 |
| 26 | Anti-rabbit IgG, HRP-linked | CST | 7074 | 1:8000 |
| 27 | Anti-mouse IgG, HRP-linked | CST | 7076 | 1:6000 |

**Supplementary Table 2.**
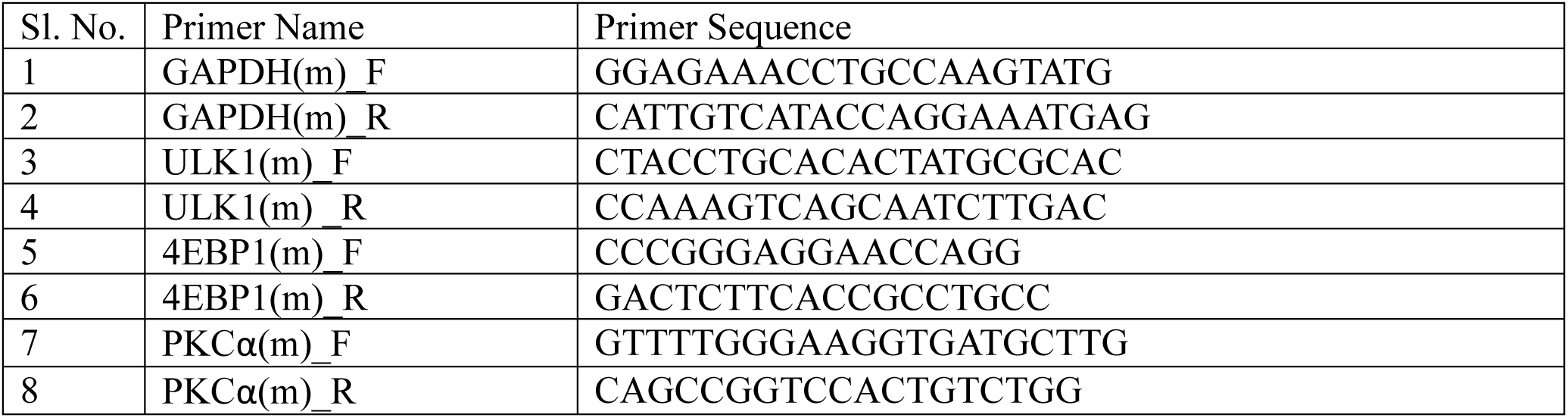
List of qPCR primers used for assessing the transcripts expression.

